# Mobility in Osteogenesis Imperfecta: A Multicenter North American Study

**DOI:** 10.1101/378190

**Authors:** Karen M. Kruger, Angela Caudill, Mercedes Rodriguez Celin, Sandesh CS Nagamani, Jay R Shapiro, Robert D Steiner, Michael B Bober, Tracy Hart, David Cuthbertson M.S., Jeff Krischer, Peter H Byers, Michaela Durigova, Francis H Glorieux, Frank Rauch, V Reid Sutton, Brendan Lee, Eric T Rush, Peter A. Smith, Gerald F. Harris

## Abstract

**Background:** Osteogenesis imperfecta (OI) is a genetic connective tissue disorder characterized by increased bone fragility and recurrent fractures. The phenotypic severity of OI has a significant influence on the ability to walk but little is known about the ambulatory characteristics, strength, or functional abilities in individuals with OI, especially in the more severe forms. To advance clinical research in OI, the Linked Clinical Research Centers, network of clinical centers in North America with significant experience in treating patients with OI, was established in 2009. The purpose of this work was to characterize mobility in OI using standard clinical assessment tools. and determine if any patient characteristics could be used to predict mobility outcomes.

**Methods:** Data were collected at five clinical sites and included age, gender, ethnicity, height, weight, use of assistive device, and bisphosphonate use and mobility metrics (age at first walk, Gillette Functional Assessment Questionnaire, Functional Mobility Scale, and distance walked in the 6 minute walk test). Linear mixed models were developed to explore the relationships between subject demographics and mobility metrics.

**Results:** The study identified 491 individuals age 3 and older. In general, the results showed minor limitations in the type I group while the more severe types showed more significant limitations in all mobility metrics analyzed. Height and weight were shown to be the most significant predictors of mobility metrics. Relationships with mobility and bisphosphonates varied with OI type and whether oral or IV was used.

**Conclusion:** This paper is the most comprehensive report of mobility in individuals with OI to date. These results are vital to understanding the mobility limitations of specific types of OI and beneficial when developing rehabilitation protocols for this population. It is important for physicians, patients, and caregivers to gain insight into severity and classification of the disease and the influence of disease-related characteristics on the prognosis for mobility.

## INTRODUCTION

Osteogenesis imperfecta (OI) is a genetic connective tissue disorder characterized by increased bone fragility and recurrent fractures [1] that has an estimated prevalence of 1:10,000 individuals [2]. OI is classified into various types depending on the disease severity and the genetic basis of the disorder. Sillence originally described four types (I- classic non-deforming, II-perinatally lethal, III-progressively deforming OI, IV-common variable OI). However, recent advances in understanding the molecular basis of OI has expanded the number of subtypes to 16 [3, 4]. Heightened diagnostic awareness and improved treatments, particularly in severe forms, has increased the number of individuals living with OI [5].

The phenotypic severity of OI has a significant influence on the ability to walk [6]. Little is known, however, about the ambulatory characteristics, strength, or functional abilities in individuals with OI, especially in the more severe forms [1]. A recent report showed children with type I OI to be as active as typically developing peers [7]. Studies using the Pediatric Outcomes Data Collection Instrument (PODCI) in OI showed little difference from control group for basic mobility and function but lower scores than their peers in sports and physical function [8]. Quantitative assessments have shown subtle a measurable deficiency of mobility and strength, mainly in type I cases [8–10]. While it is known that individuals with the severe types of OI have significant limitations in mobility, the variability within these groups has precluded any conclusions about the effects of modern treatment such as bisphosphonate medication, surgery, and physical therapy on this parameter [11, 12].

It is well-recognized that collaborative multicenter studies in rare diseases can lead to a better understanding of the natural history of disease, generate hypotheses for further research, and provide better therapeutic options for patients [13]. Providing evidence-based answers to clinically relevant questions in OI is challenged by the rarity of the condition. To advance clinical research in OI, the Linked Clinical Research Centers (LCRC), a network of five clinical centers in North America with significant experience in treating patients with OI, was established in 2009 [14, 15]. The LCRC conducted “Longitudinal Study of Osteogenesis Imperfecta”, an observational study wherein data were collected systematically across all participating sites. The purpose of this work was to characterize mobility in OI using standard clinical assessment tools. The specific goals of this paper were to report the range of various mobility metrics observed in each type of OI and determine if any patient characteristics could be used to predict mobility outcomes.

## METHODS

The LCRC is comprised of five clinical sites across North America and a data collection and analysis center [15]. The establishment of the LCRC and the subjects enrolled in the study have been previously described and followed IRB approved protocols [15]. Mobility data was collected in accordance with detailed instructions in the Manual of Operations and the quality of data was assessed at the entry point using on-line case report forms.

The following data were collected from all participants: age, gender, ethnicity, height, weight, use of assistive device, and bisphosphonate use. The mobility metrics analyzed included age at first walk (self-reported), Gillette Functional Assessment Questionnaire (FAQ) [16], Functional Mobility Scale (FMS) [17], and distance walked in the 6 minute walk test (6MWT) [18]. Data were analyzed from individuals over 3 years.

### Statistical Analysis

Multi regression analysis was used to determine the effect of predictor variables on mobility metrics. Mobility metrics analyzed were the FMS and 6MWT. Predictor variables included OI type, age at assessment, use of oral and IV bisphosphonate, height, weight, and gender. Using the preliminary results to identify predictor variables that correlated with FMS and 6MWT scores, linear mixed models were developed. Due to the pattern of interactions with OI type observed during preliminary analysis, separate models were developed for types I, III, and IV. These models were adjusted for within-individual correlations. Correlations between each mobility metric and predictor variable were examined by simple and multiple regression analyses. P values < 0.01 were considered statistically significant.

## RESULTS

From 558 individuals enrolled in the study, 491 individuals were 3 years or older (average age: 19.0 years ± 14.6 years, range 3.1 to 67.3 years). Subjects were identified with six OI types: 219 type I, 87 type III, 141 type IV, 17 type V, 10 type VI, 4 type VII, and 13 unclassified. Full subject demographic information including gender, race, age, height, and weight is included in Table 1. The age at first walk revealed that on average, walking in children with type I was delayed about 3 months compared to typically developing children (Figure 1) [19]. Individuals with type III who eventually walked were delayed 33 months, and on average, did not walk until age 3.8 years.

**Figure 1:**
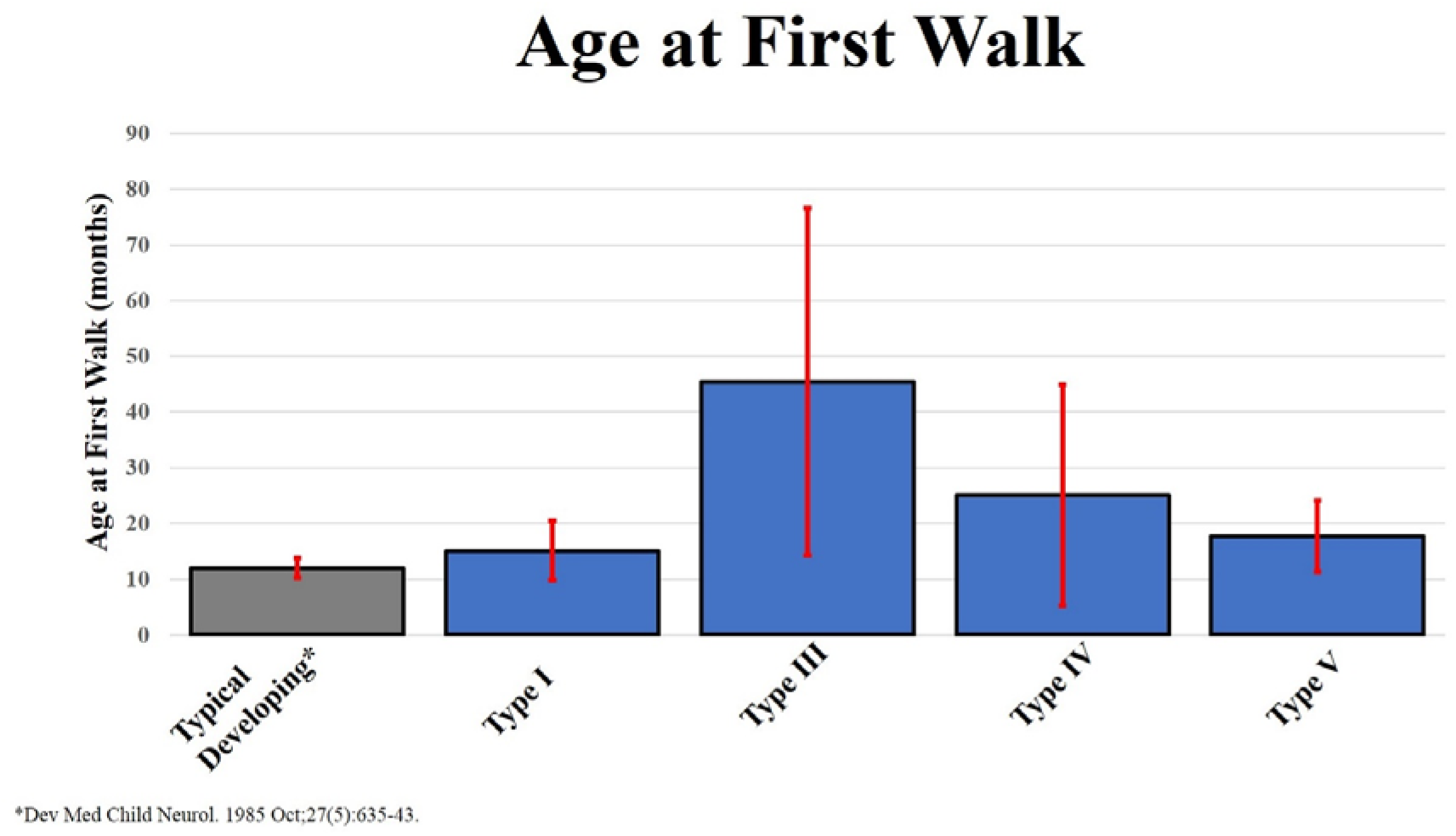
Age at First Walk (months)

**Table 1:** Demographic characteristics of patients at enrollment.

|  |  | *OI Type* |  |  |  |  |  |  |
| --- | --- | --- | --- | --- | --- | --- | --- | --- |
| *Enrollment*<br>(n=) |  | **I** | **III** | **IV** | **V** | **VI** | **VII** | **Unclassified** |
|  | Male | 104 | 34 | 63 | 6 | 5 | 0 | 7 |
|  | Female | 115 | 53 | 78 | 11 | 5 | 4 | 6 |
|  | Total | 219 | 87 | 141 | 17 | 10 | 4 | 13 |
| *Race*<br>% | White | 93.6% | 81.4% | 85.7% | 81.3% | 70.0% | 50.0% | 76.9% |
|  | Black | 0.9% | 4.7% | 6.4% | 6.3% | 0.0% | 0.0% | 15.4% |
|  | Other | 5.5% | 14.0% | 7.9% | 12.5% | 30.0% | 50.0% | 7.7% |
| *Age(yrs)* | Ave (Range) | 21.5(3.1-67.3) | 17.6(3.2-54.1) | 17.3(3.1-62.7) | 17.5(3.1-39.7) | 13.9(3.6-31.5) | 9.3(4.0-20.2) | 13.7(6.9-32.9) |
| *Height*<br>(cm) | Ave (Range) | 150.5(55.9-188) | 100.9(48.4-141) | 133.448(70.3-170.1) | 132(78.2-177.5) | 132.5(106.0-162.4) | 112.4(76.7-98) | 122.2(98-152.7) |
| *Weight (kg)* | Ave (Range) | 53.9(12.5-168.0) | 29.5(5.9-98.0) | 46.3(10.8-111.4) | 53.8(12.5-160.1) | 42.7(22.9-64.8) | 38.3(13.0-85.5) | 34.7(17.6-54.8) |

The FAQ walking ability scores revealed individuals with type I OI had the highest scores (average score 9.6, range 5-10, Figure 2). Individuals with type IV OI had lower scores (average 6.7, range 1-10), while individuals with type III OI scored 4.1 on average (range 1-10). Results assessing for limitations to walking varied depending on OI type. Pain concerns ranked highest for type I (29%), weakness highest for type III (43%), while safety ranked highest for type IV (40%) (Figure 3).

**Figure 2:**
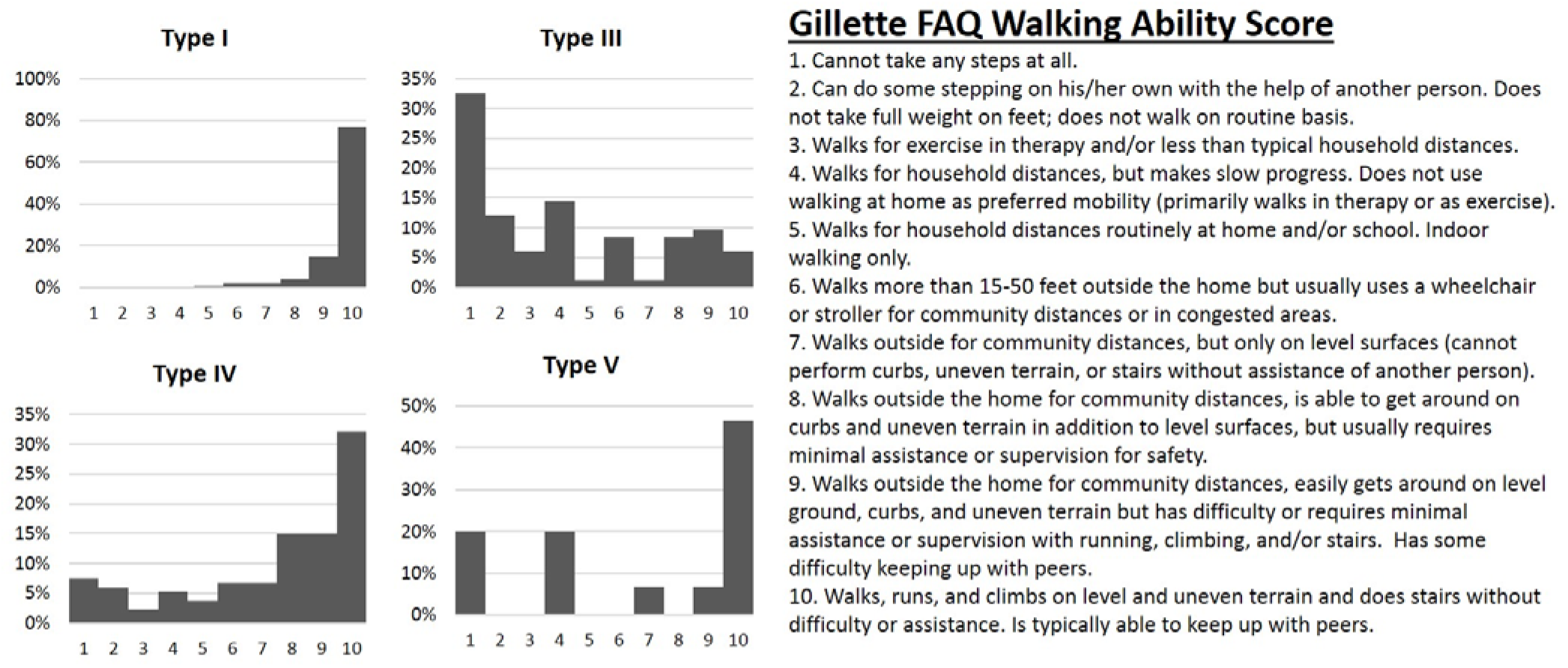
Gillette Walking Ability Score

**Figure 3:**
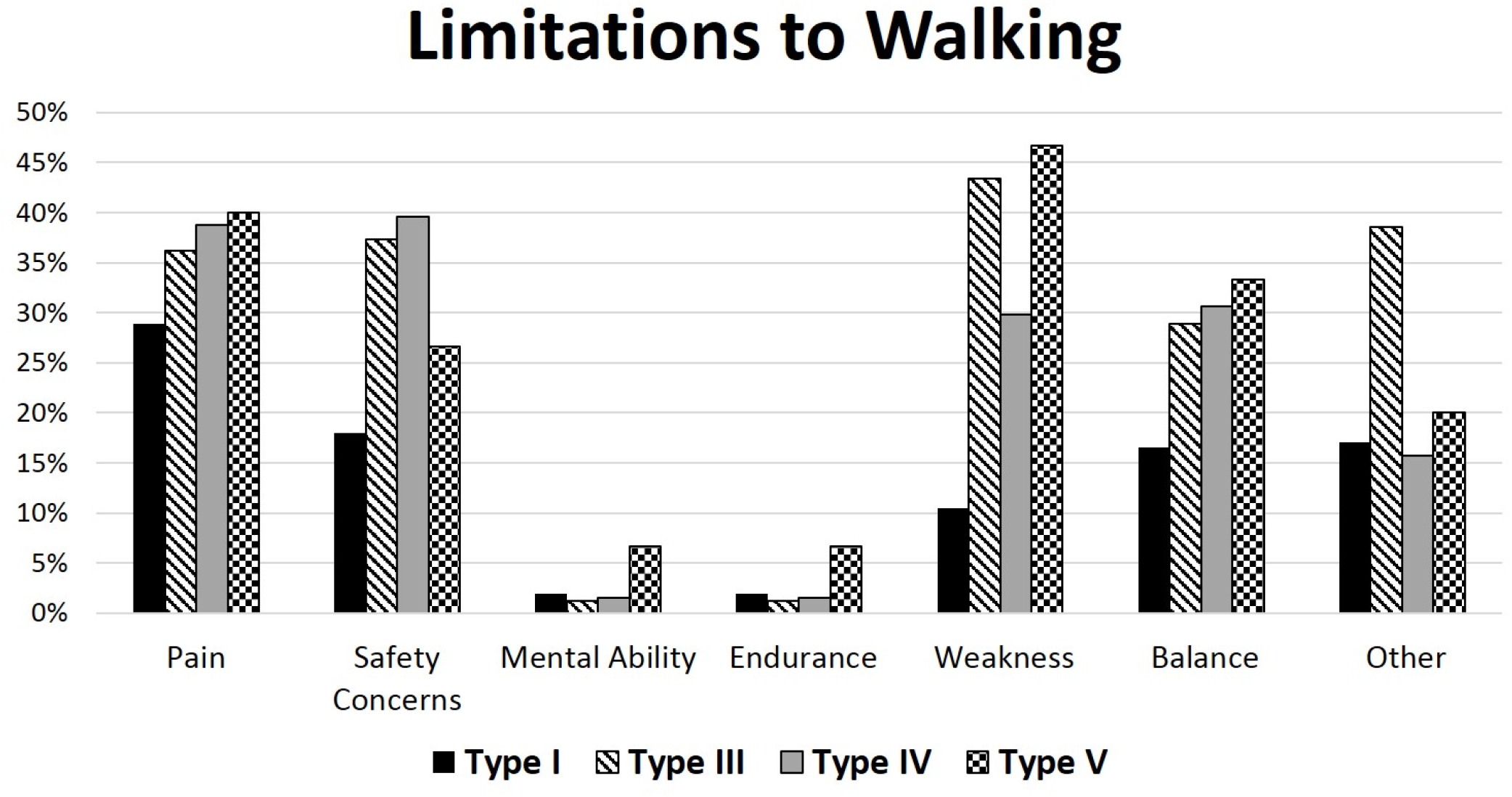
Limitations to Walking

Clinician-assigned FMS scale results demonstrated that most of individuals with type I OI could walk completely independently on all surfaces while those with type III OI required a wheelchair or walker, especially at longer distances (Figure 4). Overall, individuals with OI of all types walked shorter distances in the 6MWT than healthy adults (Figure 5). On average, the distances walked by individuals with type I OI were 30% lower than typically developing while those with type III who were able to complete the test had an average decrease of 62%.

**Figure 4:**
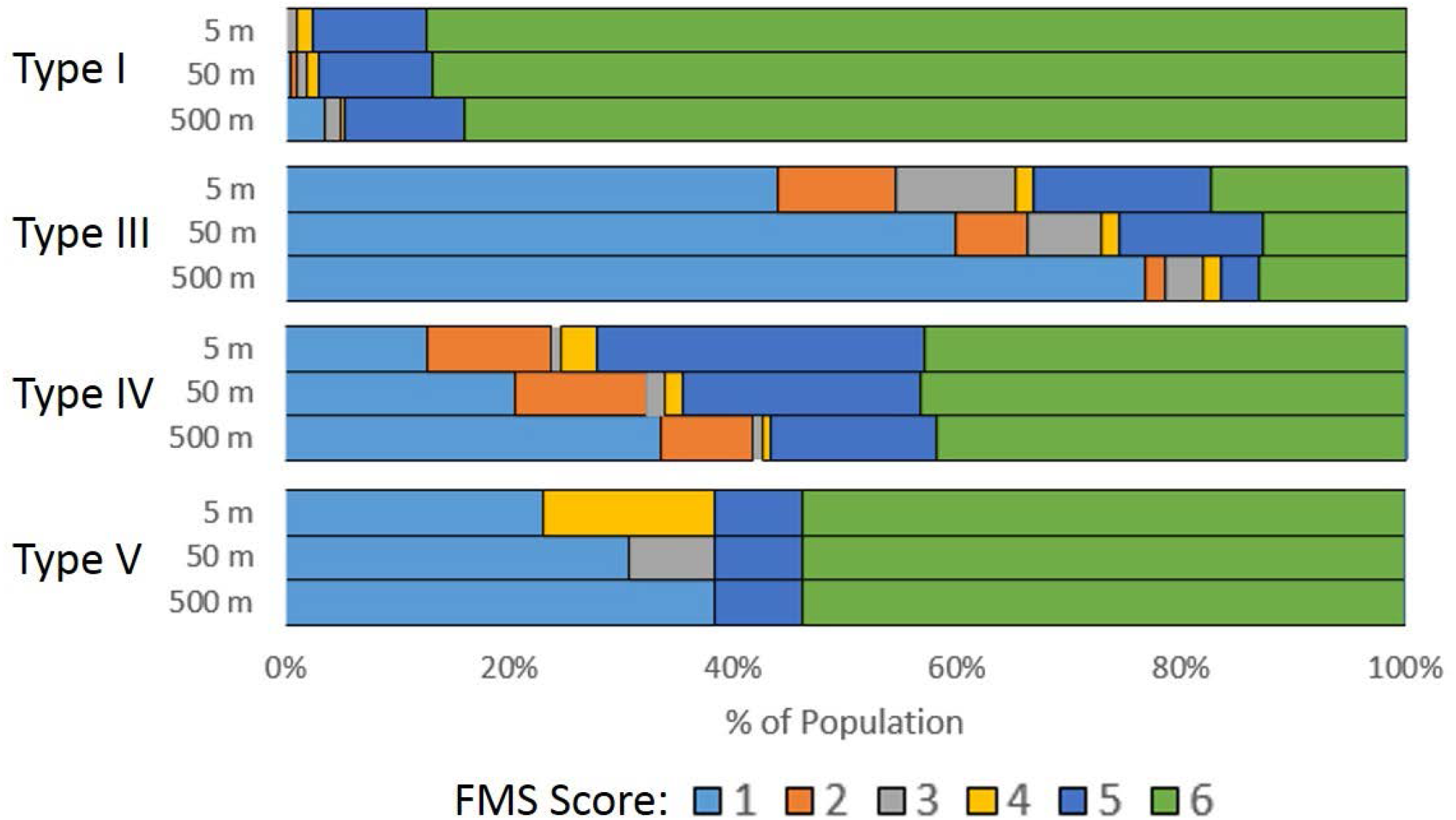
Functional Mobility Scale

**Figure 5:**
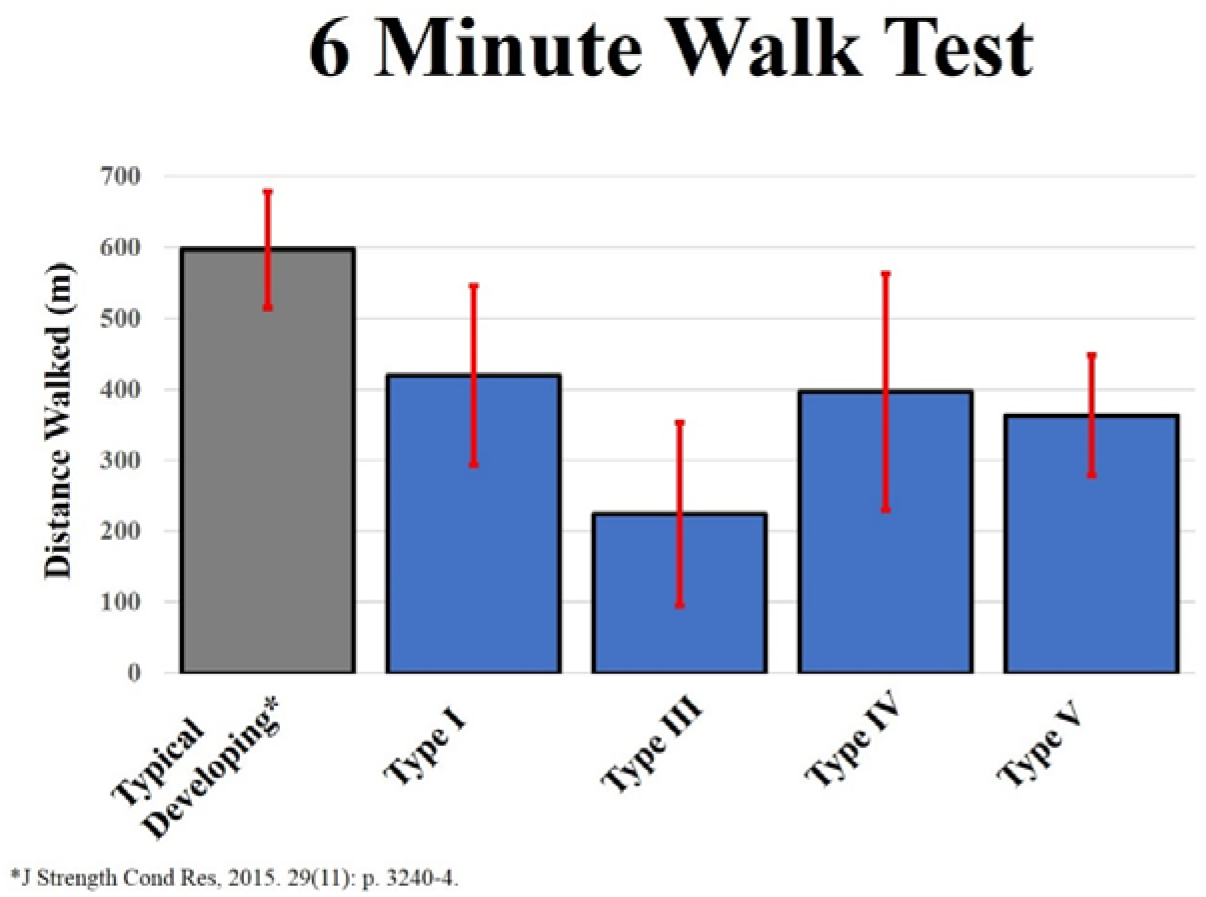
Distance walked in 6 Minute Walk Test (m)

The FAQ 22 questionnaire (Table 2) revealed that the majority of individuals with type I OI could perform tasks such as walking while carrying an object (99%), walk up and down stairs using the railing (99%), and taking a step backwards (98%). In contrast, only 25-40% of individuals with type III OI could complete these tasks while 69-77% of individuals with type IV OI could. Ice/roller skating was ranked as the lowest task completion rate with 33%, 2%, and 10% of individuals of OI types I, III, and IV, respectively, reporting being able to complete the task.

**Table 2:** Gillette FAQ Tasks. Number indicate the percentage of population responding “yes” to being able to complete each task.

| **Task** | **OI Type** |  |  |  |  |  |
| --- | --- | --- | --- | --- | --- | --- |
|  | **I** | **III** | **IV** | **V** | **VI** | **VII** |
| Can maneuver in tight areas | 97% | 36% | 78% | 73% | 50% | 50% |
| Can take steps backward | 98% | 43% | 74% | 73% | 60% | 50% |
| Get on and off a bus by him/herself | 89% | 13% | 49% | 53% | 30% | 50% |
| Hop on left foot | 77% | 10% | 37% | 27% | 20% | 25% |
| Hop on right foot | 78% | 10% | 36% | 27% | 10% | 25% |
| Ice skate or roller skate | 33% | 2% | 10% | 7% | 0% | 0% |
| Jump rope | 63% | 5% | 24% | 20% | 10% | 25% |
| Jumps off a single step | 81% | 13% | 39% | 33% | 10% | 25% |
| Kick a ball with left foot | 92% | 42% | 71% | 60% | 70% | 50% |
| Kick a ball with right foot | 91% | 47% | 75% | 67% | 70% | 50% |
| Ride 2 wheel bike | 66% | 4% | 25% | 20% | 0% | 25% |
| Ride 3 wheel bike | 63% | 36% | 52% | 27% | 40% | 50% |
| Ride an escalator | 85% | 10% | 49% | 47% | 40% | 0% |
| Runs well | 76% | 6% | 37% | 20% | 10% | 25% |
| Step over an object, right foot first | 96% | 43% | 72% | 67% | 60% | 50% |
| Step over an object, left foot first | 96% | 43% | 74% | 60% | 50% | 50% |
| Steps up and down curb | 97% | 23% | 63% | 53% | 60% | 50% |
| Walk carrying a fragile object | 98% | 25% | 69% | 73% | 60% | 50% |
| Walk carrying an object | 99% | 40% | 77% | 73% | 60% | 50% |
| Walk up and down stairs using the railing | 99% | 30% | 75% | 67% | 60% | 50% |
| Walk up and down stairs using railing | 83% | 7% | 42% | 33% | 30% | 25% |

Requirement of assistive devices was variable by OI type (Table 3). Of the individuals that responded, 4%, 40%, and 24% of those with OI types I, III, and IV, respectively, required assistive devices. A walker was the most commonly used device, followed crutches and a cane. Bisphosphonate use was reported by just over half of type I individuals while a large majority of individuals with the other types received either oral and IV bisphosphonate (Table 4).

**Table 3:** Use of Assistive Devices

|  | *I* | *III* | *IV* | *V* | *VI* | *VII* | *Unclassified* |
| --- | --- | --- | --- | --- | --- | --- | --- |
| *Yes* | 4% | 40% | 24% | 25% | 50% | 50% | 17% |
| *No* | 96% | 60% | 76% | 75% | 50% | 50% | 83% |
| *Walker* | 4 | 24 | 16 | 2 | 4 | 1 | 1 |
| *Cane* | 2 | 2 | 6 | 1 | 1 | 1 |  |
| *Crutches* | 3 | 7 | 10 | 1 |  |  | 1 |
| *Unknown* |  | 1 | 2 |  |  |  |  |

**Table 4:** History of Bisphosphonate Use.

| **Type** | **None** | **Oral** | **IV** |
| --- | --- | --- | --- |
| **I** | 45% | 19% | 35% |
| **III** | 10% | 13% | 77% |
| **IV** | 13% | 15% | 72% |
| **V** | 33% | 7% | 60% |
| **VI** | 0% | 9% | 91% |
| **VII** | 0% | 0% | 100% |

The results of the bivariate analysis showed that when looking at one predictor variable, weight, height, use of IV bisphosphonate, and OI type were significant for all mobility metrics assessed. Age and gender were not significant predictors for any of the mobility metrics analyzed. Multi regression analysis showed fewer correlations of statistical significance (Table 7). The results of this analysis can be used to predict these mobility outcomes for an individual subject. For example,

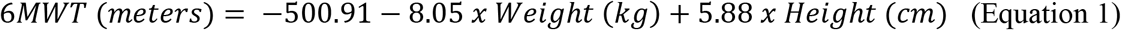

**Table 7:** Results of Regression Analysis. * indicates variable was a statistically significant predictor of mobility metric. Values represent intercept and slope for multi-variable regression equations as reported in Equation 1.

|  | 6MWT | FMS - 5 m | FMS - 50 m | FMS - 500 m |
| --- | --- | --- | --- | --- |
| (Intercept) | -500.907 | -4.973 | -0.974 | 1.149 |
| Age | 3.367 | 0.009 | -0.002 | 0.003 |
| Weight | -8.053* | -0.019 | 0.009 | -0.005 |
| Height | 5.881* | 0.074* | 0.047* | 0.044 |
| Years (IV Bisphosphonate) | -10.623 | 0.024 | 0.046 | 0.054 |
| Years (Oral Bisphosphonate) | -5.917 | -0.048 | -0.06 | -0.068 |

Many interactions were observed between the variables analyzed; therefore, the final interaction analysis was performed by OI type. All groups except type I individuals who were not given bisphosphonate showed increased 6MWT distances with increasing height (Figure 6). This trend did not continue when analyzing correlation with weight (Figure 7). The type III group showed a strong correlation between increasing weight and increasing 6MWT for individuals on either IV or no bisphosphonates while the type I group showed only a slight trend of increased 6MWT distance with increased weight for those given IV bisphosphonate.

**Figure 6:**
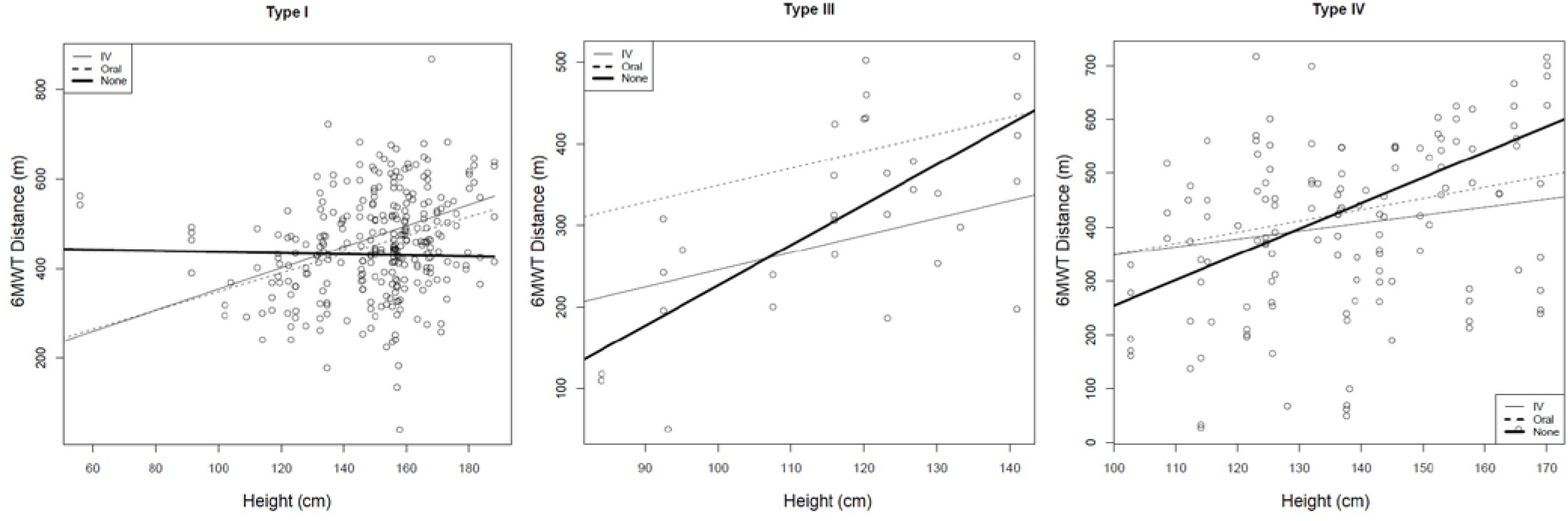
Relationship between height and 6MWT by OI Type

**Figure 7:**
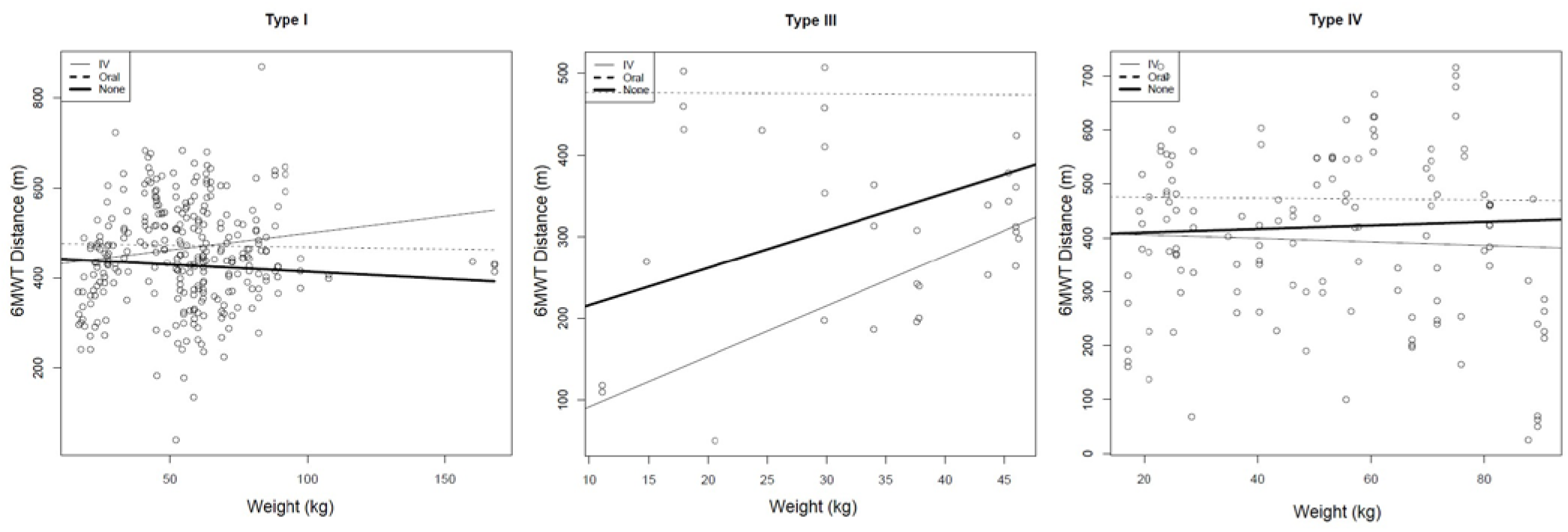
Relationship between weight and 6MWT by OI Type

## DISCUSSION

The prognosis for mobility of a child with OI is of clinical interest when setting goals for rehabilitation. Understanding disease related characteristics specific to each type allows for more accurate patient comparisons and expectations. Children and young adults with OI often experience a reoccurring cyclical pattern involving fracture, weakness, and deconditioning which leads to functional limitations [9]. Maintaining the highest level of physical activity possible without injury is a primary goal when setting rehabilitation strategies. Successful rehabilitative intervention in mild to moderate OI seeks to maintain an appropriate loading environment to promote independence in the face of impairments from bone fragility.

Physical activity is important in maintaining strength and preventing other disorders including cardiovascular diseases and diabetes [20]. Furthermore, youth with OI expressed disappointment in missing out on physical activities such as running due to tiring more easily than their more able bodied peers [21]. Therefore, all efforts should be made to maximize mobility in individuals with OI. Efforts have been made to study the impact of functional exercise on children with OI and have found that a standardized fitness program can improve aerobic performance and muscle strength [22]. Several reports have also been published on the increased prevalence of obesity in patients with OI with type III patients being the most severely affected [23–25] further demonstrating the importance of promoting physical activity within this population.

The results of the statistical analysis showed only relatively minor associations between the predictor variables and mobility outcome measures. As is shown in our results, height deficit is related to worse mobility outcomes, agreeing with literature where height has been related with severity of the condition [26, 27]. While BMI was not explicitly used as a predictor variable for this analysis, the interactions of height and weight were explored as part of the multivariate analysis and showed no associations for any of the mobility outcomes analyzed. These results indicate that the predictor variables commonly used to describe individuals with OI are not adequate to be able to prescribe treatment. Future work with this consortium and other groups who treat individuals with OI should use more detailed data including strength assessment and gait analysis to assess and set rehabilitation goals. There have been some reports on strength and gait characteristics which had small sample sizes and were limited to type I cases [8, 9, 28, 29]. Future work should include larger patient populations of all OI types to gain further understanding of functional limitations associated with various types of OI. This information will be vital to prescribing the best treatment to address these limitations. Normative data presented here such as 6MWT and other parameters can also be used as outcome measures in clinical trials in this population.

How to prescribe treatment remains an important clinical question. Zeitlin et al. have stressed treatment for OI, particularly in severe cases, needs to be a multi-disciplinary approach including bisphosphonate therapy, corrective orthopedic interventions, and physical therapy [11]. The criteria for dividing individuals with OI into treatment groups has been subject to debate. Engelbert et al suggested degree of intervention needed depends on severity of the clinical phenotype [12]. In contrast, Sousa suggested the Silence classification does not direct clinical patient care regarding treatments and interventions utilized and that treatment criteria should be established based on the number of fractures experienced per year [30]. They argued that the Silence classification is not uniformly predictive of which patients with OI will qualify for pamidronate therapy. Mobility outcomes and height have also been suggested as features to be taken in account for severity assessment and clinical classification of OI [31].

Benefits of bisphosphonates in mobility remain inconclusive [32]. Land et al reported that cyclical pamidronate treatment improves mobility, ambulation level, and muscle force in children with moderate to severe OI [33] while Seikaly reported improvements in well-being scores and self-care abilities but no change in mobility in patients treated with alendronate [34]. Other reports found no changes in mobility or functional outcomes using oral [35–37] or intravenous bisphosphonates [38]. In agreement with these studies, our results showed no significant relationship between years on bisphosphonate treatment and mobility outcomes. However, it is important to note that the effect of the medication is affected by different factors such as age of starting, duration of the treatment, severity of the condition, and the association with other multidisciplinary approaches including physiotherapy and corrective surgery [39–41]. The most important predictor of mobility remains phenotype.

This work is limited in that the FAQ and FMS are subjective in nature. Both the 10 point walking scale [16] and 22-item skill set questionnaire [42] have been validated as suitable for measuring locomotor skill ability in children. While they provide a better overall view of community functional ability, they are subjective to a degree and quantitative kinematics would be a useful tool to use in conjunction with these data. Some individuals with more severe forms of OI scored higher than expected on several aspects of the FAQ. One possible explanation for this is that children with OI types III and IV sometimes use wheelchairs for sports activities and have different expectations than unaffected children [43]. This makes comparison of patient reported outcome in these heterogeneous groups difficult.

The FMS, while still subjective, is a clinician assigned score which could be reasonably expected to yield more consistent results than the FAQ while 6MWT is an objective measure. Neither of these had a strong correlation to the patient characteristics analyzed. This suggests more detailed mobility data such as strength testing and gait analysis should be analyzed to gain insight into mobility limitations. This has been analyzed in small groups of individuals with type I OI. Strength testing revealed decreased heel-rise strength and ankle isometric plantar flexion strength [9]. Results of a gait study showed the OI population demonstrated abnormal gait parameters including increased double support, delayed foot off, reduced ankle range of motion and plantarflexion during third rocker, along with greater ankle power absorption during terminal stance and reduced ankle power generation during push off [8, 29]. Together, these findings provide a comprehensive description of gait characteristics among a group of individuals with type I OI. Such data inform clinicians about specific gait deviations in this population allowing clinicians to recommend more focused interventions however no such analysis has been completed on the more severe OI types.

In conclusion, this study reports cross-sectional data from the largest study of mobility OI conducted to date. Results are vital to understanding the mobility limitations of specific types of OI and beneficial when developing rehabilitation protocols for this population. It is important for physicians, patients, and caregivers to gain insight into severity and classification of the disease and the influence of disease-related characteristics on the prognosis for mobility.

## Acknowledgments

We would like to thank Dr. Sergey Tarima for statistical consultation and the members of the Brittle Bone Disease (BBD) Consortium (Jean Marc Retrouvey, Faculty of Dentistry, McGill University, Montreal; Paul Esposito, University of Nebraska Medical Center, Omaha; David Eyre, Department of Orthopedic and Sports Medicine, University of Washington, Seattle; Danielle Gomez, Shriners Hospital for Children, Tampa; Gerald Harris, Marquette University and Medical College of Wisconsin; Mahim Jain, Departments of Bone and Osteogenesis Imperfecta, Kennedy Krieger Institute, Baltimore; Deborah Krakow, Departments of Orthopedic Surgery and Obstetrics and Gynecology, David Geffen School of Medicine, University of California, Los Angeles; Eric Orwoll, Department of Medicine, Division of Endocrinology, Oregon Health & Science University, Portland; Cathleen Raggio, Hospital for Special Surgery, New York; Laura Tosi, Bone Health Program, Children’s National Health System, Washington, DC). Funding was provided by Children’s Brittle Bone Foundation, OI Foundation, and NIDLRR Grant 90AR5022-01.

